# Chromatinization of *Escherichia coli* with archaeal histones

**DOI:** 10.1101/660035

**Authors:** Maria Rojec, Antoine Hocher, Matthias Merkenschlager, Tobias Warnecke

## Abstract

Nucleosomes restrict DNA accessibility throughout eukaryotic genomes, with repercussions for replication, transcription, and other DNA-templated processes. How this globally restrictive organization emerged from a presumably more open ancestral state remains poorly understood. Here, to better understand the challenges associated with establishing globally restrictive chromatin, we express histones in a naïve bacterial system that has not evolved to deal with nucleosomal structures: *Escherichia coli*. We find that histone proteins from the archaeon *Methanothermus fervidus* assemble on the *E. coli* chromosome *in vivo* and protect DNA from micrococcal nuclease digestion, allowing us to map binding footprints genome-wide. We provide evidence that nucleosome occupancy along the *E. coli* genome tracks intrinsic sequence preferences but is disturbed by ongoing transcription and replication. Notably, we show that higher nucleosome occupancy at promoters and across gene bodies is associated with lower transcript levels, consistent with local repressive effects. Surprisingly, however, this sudden enforced chromatinization has only mild repercussions for growth, suggesting that histones can become established as ubiquitous chromatin proteins without interfering critically with key DNA-templated processes. Our results have implications for the evolvability of transcriptional ground states and highlight chromatinization by archaeal histones as a potential avenue for controlling genome accessibility in synthetic prokaryotic systems.

## INTRODUCTION

All cellular systems face the dual challenge of protecting and compacting their resident genomes while making the underlying genetic information dynamically accessible. In eukaryotes, this challenge is solved, at a fundamental level, by nucleosomes, ~147bp of DNA wrapped around an octameric histone complex. Nucleosomes can act as platforms for the recruitment of transcriptional silencing factors such as heterochromatin protein 1 (HP1) in animals (Danzer, 2004; Zhao *et al*, 2000) and Sir proteins in yeast (Gartenberg & Smith, 2016), but can also directly render binding sites inaccessible to transcription factors (Beato & Eisfeld, 1997; Zhu *et al*, 2018). As a consequence, gene expression in eukaryotes is often dependent on the recruitment of chromatin remodelers. By controlling access to DNA, histones play a key role in maintaining a low basal rate of transcription in eukaryotic cells and have therefore been described as the principal building blocks of a restrictive transcriptional ground state (Struhl, 1999).

Histones are not confined to eukaryotes, but also common in archaea (Adam *et al*, 2017; Henneman *et al*, 2018). They share the same core histone fold but lack N-terminal tails, which are the prime targets for posttranslational modifications in eukaryotes (Henneman *et al*, 2018). They form dimers in solution and bind DNA as tetrameric complexes that wrap ~60bp instead of ~147bp of DNA (Reeve *et al*, 2004). At least in some archaea, these tetrameric complexes can be extended, in dimer steps, to form longer oligomers that wrap correspondingly more DNA (~90bp, ~120bp, etc.) and assemble without the need for dedicated histone chaperones (Xie & Reeve, 2004; Mattiroli *et al*, 2017; Maruyama *et al*, 2013). Archaeal and eukaryotic nucleosomes preferentially assemble on DNA that is more bendable, a property associated, on average, with elevated GC content and the presence of certain periodically spaced dinucleotides, notably including AA/TT (Ammar *et al*, 2011; Nalabothula *et al*, 2013; Pereira & Reeve, 1999; Bailey *et al*, 2000; 2002; Ioshikhes *et al*, 2011). They also exhibit similar positioning around transcriptional start sites (Ammar *et al*, 2011; Nalabothula *et al*, 2013), which are typically depleted of nucleosomes and therefore remain accessible to the core transcription machinery. Whether archaeal histones play a global restrictive role akin to their eukaryotic counterparts, however, remains poorly understood, as does their involvement in transcription regulation more generally (Gehring *et al*, 2016).

Thinking about the evolution of restrictive chromatin and its molecular underpinnings, we wondered how the presence of histones would affect a system that is normally devoid of nucleosomal structures. How would a cell that has neither dedicated nucleosome remodelers nor co-evolved sequence context cope with chromatinization? Could global chromatinization occur without fundamentally interfering with DNA-templated processes? How easy or hard is it to transition from a system without histones to one where histones are abundant?

Motivated by these questions, we built *Escherichia coli* strains expressing histones from the hyperthermophilic archaeon *Methanothermus fervidus* (HMfA or HMfB), on which, thanks to the pioneering work of Reeve and co-workers, much of our foundational knowledge about archaeal histones is based. HMfA and HMfB are 85% identical at the amino acid level but differ with regard to their DNA binding affinity and expression across the *M. fervidus* growth cycle, with HMfB more prominent towards the latter stages of growth and able to provide greater DNA compaction *in vitro* (Sandman *et al*, 1994; Marc *et al*, 2002). We find that HMfA and HMfB, heterologously expressed in *E. coli*, bind to the *E. coli* genome and protect it from micrococcal nuclease (MNase) digestion, allowing us to map nucleosomes in *E. coli in vivo*. We present evidence for sequence-dependent nucleosome positioning and occupancy and consider how the binding of histones affects transcription on a genome-wide scale. Importantly, we find evidence for local repressive effects associated with histone occupancy yet only mild repercussions for growth and cell morphology. Overall, *E. coli* copes remarkably well with enforced chromatinization, supporting the notion that archaeal histones are leaky barriers when it comes to inhibiting transcription. Our findings have implications for how histones became established as global repressive regulators during the evolution of eukaryotes, highlight the utility of heterologous systems in understanding major evolutionary transitions, and might inform future efforts to engineer restrictive chromatin in synthetic prokaryotic systems.

## RESULTS

### *Archaeal histones bind the* E. coli *genome* in vivo, *assemble into oligomers, and confer protection from MNase digestion*

We transformed an *E. coli* K-12 MG1655 strain with plasmids carrying either *hmfA* or *hmfB*, codon-optimised for expression in *E. coli* and under the control of a rhamnose-inducible promoter (see Methods, Figure S1). Below, we will refer to these strains as Ec-hmfA and Ec-hmfB, respectively, with Ec-EV being the empty vector control strain (Table S1). Following induction, both histones are expressed at detectable levels and predominantly found in the soluble fraction of the lysate in both exponential and stationary phase (Figure S2). We did not observe increased formation of inclusion bodies. Based on dilution series with purified histones (see Methods, Figure S2), we estimate HMfA:DNA mass ratios of up to ~0.6:1 in exponential and ~0.7:1 in stationary phase, which corresponds to 1 histone tetramer for every 76bp (64bp) in the *E. coli* genome. Given that a tetramer wraps ~60bp of DNA, this implies a supply of histones that is, in principle, sufficient to cover most of the *E. coli* genome. However, it is important to note that, at any given time, not all histones need to be associated with DNA.

We carried out MNase digestion experiments using samples extracted in late exponential and stationary phase, corresponding to 2h and 16-17h after induction, respectively (see Methods). In response to a wide range of enzyme concentrations, MNase digestion of fixed chromatin from Ec-hmfA/B (see Methods) yields a ladder-like pattern of protection that is not observed in Ec-EV (Figure 1A-B). Across many replicates, we could usually discriminate the first four rungs of the ladder, with the largest rung at 150bp. On occasion, we observe multiple larger bands (e.g. for Ec-hmfA in Figure 1A). Sequencing digestion fragments <160bp using single-end Illumina technology recapitulates the read length distribution seen on gels, with peaks around 60bp, 90bp, 120bp, and 150bp (Figure 1C), consistent with oligomerization dynamics described for archaeal histones in their native context (Maruyama *et al*, 2013; Mattiroli *et al*, 2017). Indeed, we obtained remarkably similar digestion profiles when we applied the same protocol, modified to account for altered lysis requirements (see Methods), to *M. fervidus* cultures (Figure 1A). Modal fragment sizes of ~60bp and ~90bp in exponential and stationary phase (Figure 1C), respectively, suggest that larger oligomers become more prevalent later in the growth cycle, which might reflect elevated histone:DNA ratios but also reduced perturbation from replication and transcription, as further discussed below. In exponential phase only, an additional peak is evident at ~30bp. Fragments of this size were previously observed during *in vitro* reconstitution experiments with HMfA/B and, at the time, attributed to the binding of histone dimers (Grayling *et al*, 1997). However, in our digestion regime this peak is also present in Ec-EV, and we cannot therefore rule out the possibility that it is caused by specifics of the digestion protocol, library construction or native *E. coli* proteins found exclusively in exponential phase. Below, we therefore focus on larger peaks (60bp, 90bp, etc.) that are absent from Ec-EV, but present in *M. fervidus* and our histone-bearing *E. coli* strains.

**Figure 1.**
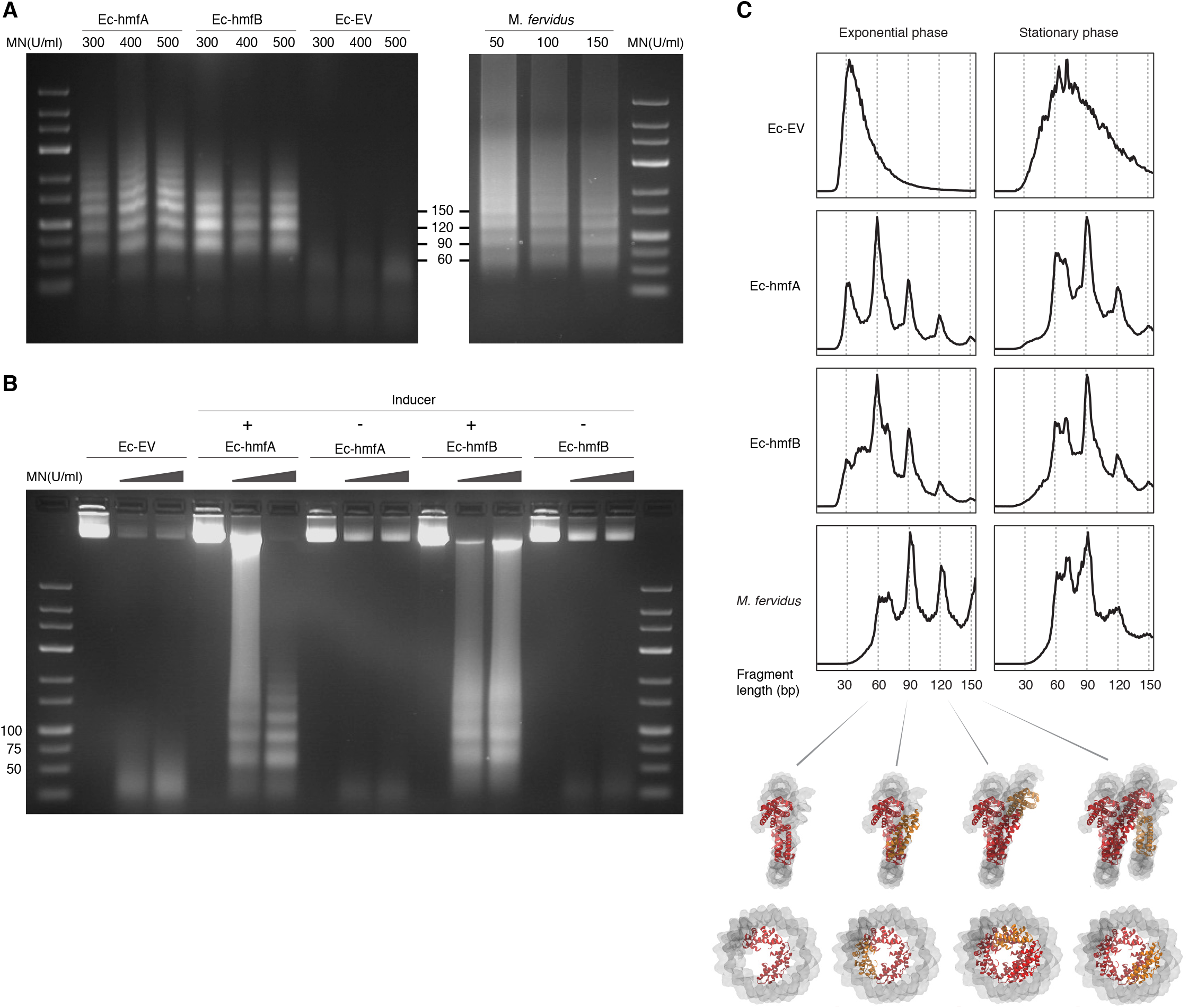
MNase digestion of *M. fervidus* and *E. coli* strains expressing *M. fervidus* histones. **A**. Agarose gel showing profiles of DNA fragments that remain protected at different MNase (MN) concentrations. **B**. Ladder-like protection profiles are only observed when *hmfA/B* expression is induced. **C.** Length distribution profiles of sequenced fragments show peaks of protection at multiples of 30bp in histone-expressing strains. Structural views below highlight how these 30bp steps would correspond to the addition or removal of histone dimers, starting from the crystal structure of a hexameric HMfB complex (PDB: 5t5k), which wraps ~90bp of DNA.

### *Intrinsic sequence preferences govern nucleosome formation along the* E. coli *genome*

Mapping digestion fragments to the *E. coli* genome, we find that binding is ubiquitous. On a coarse scale, coverage across the chromosome appears relatively even (Figure 2A). On a more local scale, however, protected fragments group into defined binding footprints (Figure 2C). Local occupancy (measured for 60bp windows, overlapping by 30bp) is highly correlated across replicates (Figure 2D), consistent with non-random binding. Ec-hmfA and Ec-hmfB are also highly correlated (Figure 2E); minor differences may reflect subtly different binding preferences, as previously reported (Bailey *et al*, 2000). Areas of apparent histone depletion often coincide with AT-rich domains (Figure 2C, F). In particular, nucleosomes are depleted from AT-rich transcriptional start sites (TSSs), mimicking a key aspect of nucleosome architecture in eukaryotes and archaea (Figure 2G).

**Figure 2.**
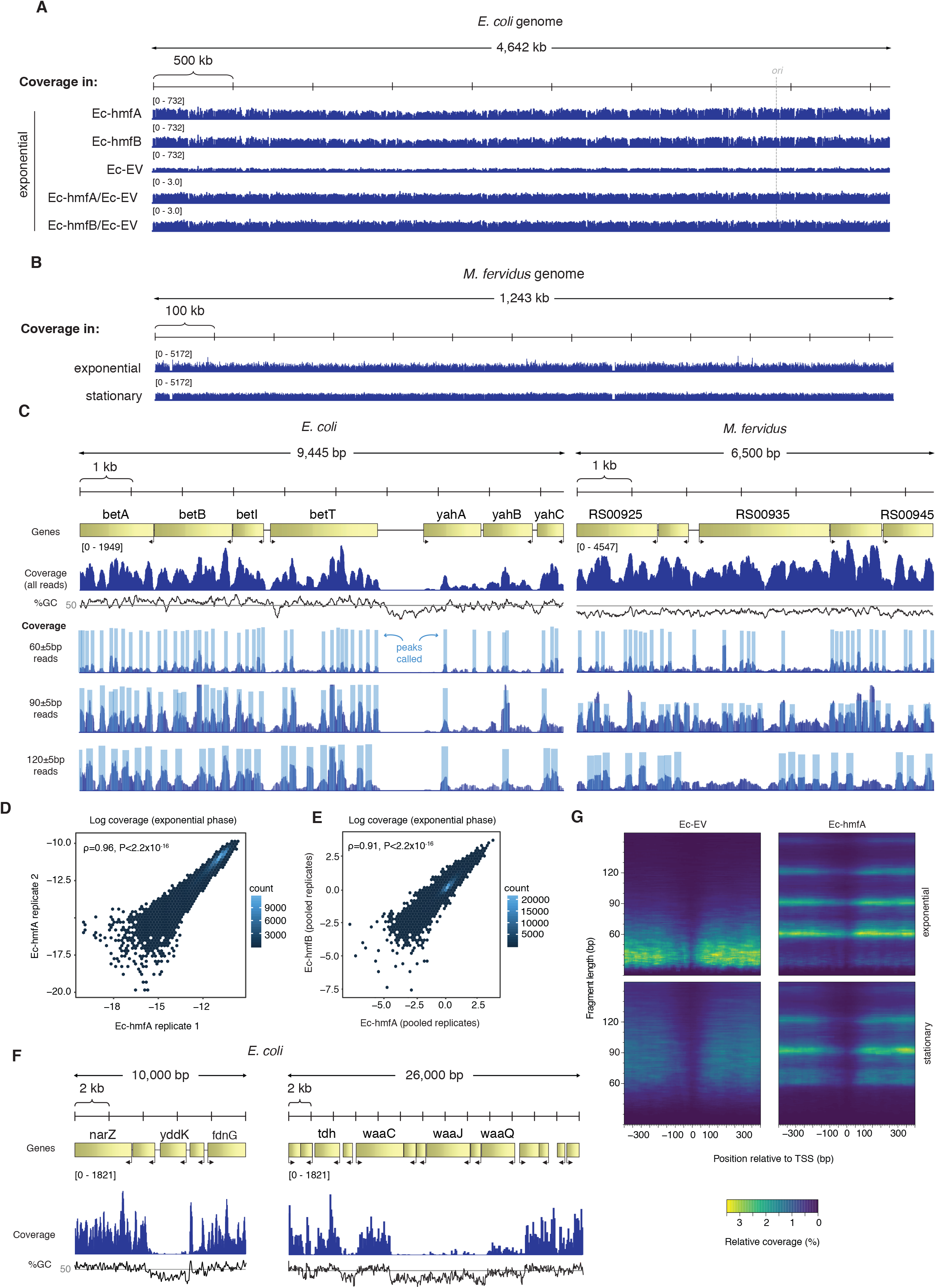
Distribution of MNase-protected fragments across the *E. coli* genome. **A.** Genome-wide coverage (and normalized coverage) tracks of MNase-protected fragments along the *E. coli* K-12 MG1655 and (**B**) the *M. fervidus* genome. **C.** Fragments of defined size cluster into footprints in *E. coli* and *M. fervidus*, as illustrated for two example regions. **D.** Correlation in coverage measured for two biological replicates of Ec-hmfA. Coverage here is expressed as a proportion of total reads in a given replicate. **E.** Correlation in normalized coverage between Ec-hmfA and Ec-hmfB. Reads were pooled across replicates for each strain. **F.** Two examples from Ec-hmfA highlighting that drops in coverage frequently correspond to regions of low GC content. **G.** Coverage as a function of both distance from experimentally defined transcriptional start sites (see Methods) and fragment size.

The above observations are consistent with a role for sequence composition in determining nucleosome positioning and/or occupancy, but likely also reflect known MNase preferences for AT-rich DNA (see Ec-EV in Figure 2G in particular). To discriminate between these two factors, we first analyzed read-internal nucleotide enrichment patterns, which should be unaffected by MNase bias. Considering fragments of exact size 60bp (90bp, etc. see Methods), we find dyad-symmetric nucleotide enrichment patterns that are absent from size-matched Ec-EV fragments but mirror what is seen in fragments from native *M. fervidus* digests (Figure 3A) (Hocher *et al*, 2019). Next, to disentangle conflated signals of MNase bias and nucleosomal sequence preferences directly, and to assess their relative impact on inferred occupancy across the genome, we normalized coverage in Ec-hmfA/B by coverage in Ec-EV (see Methods). We then trained LASSO models for different fragment size classes (60bp, 90bp, 120bp) to predict normalized occupancy across the genome from the underlying sequence, considering all mono-, di-, tri-, and tetra-nucleotides as potential predictive features (see Methods, Table S2). We find that sequence is a good predictor of normalized occupancy in stationary phase (Figure 3B-C), particularly for larger fragments (e.g. 120bp footprints in Ec-hmfA: ρ=0.72, P<2.2×10^−16^; 120bp footprints in Ec-hmfB: ρ=0.76, P=<2.2×10^−16^, Figure 3C), with simple GC content capturing much of the variability in occupancy (Figure 3B, D). Interestingly, however, the predictive power of sequence is dramatically reduced in exponential phase (Figure 3B, D). While regions of low GC content still exhibit low histone occupancy, as evident in the left-hand tails of the distributions in Figure 3D, the relationship breaks down for the bulk of the genome that has medium to high GC content. Why would this be?

**Figure 3.**
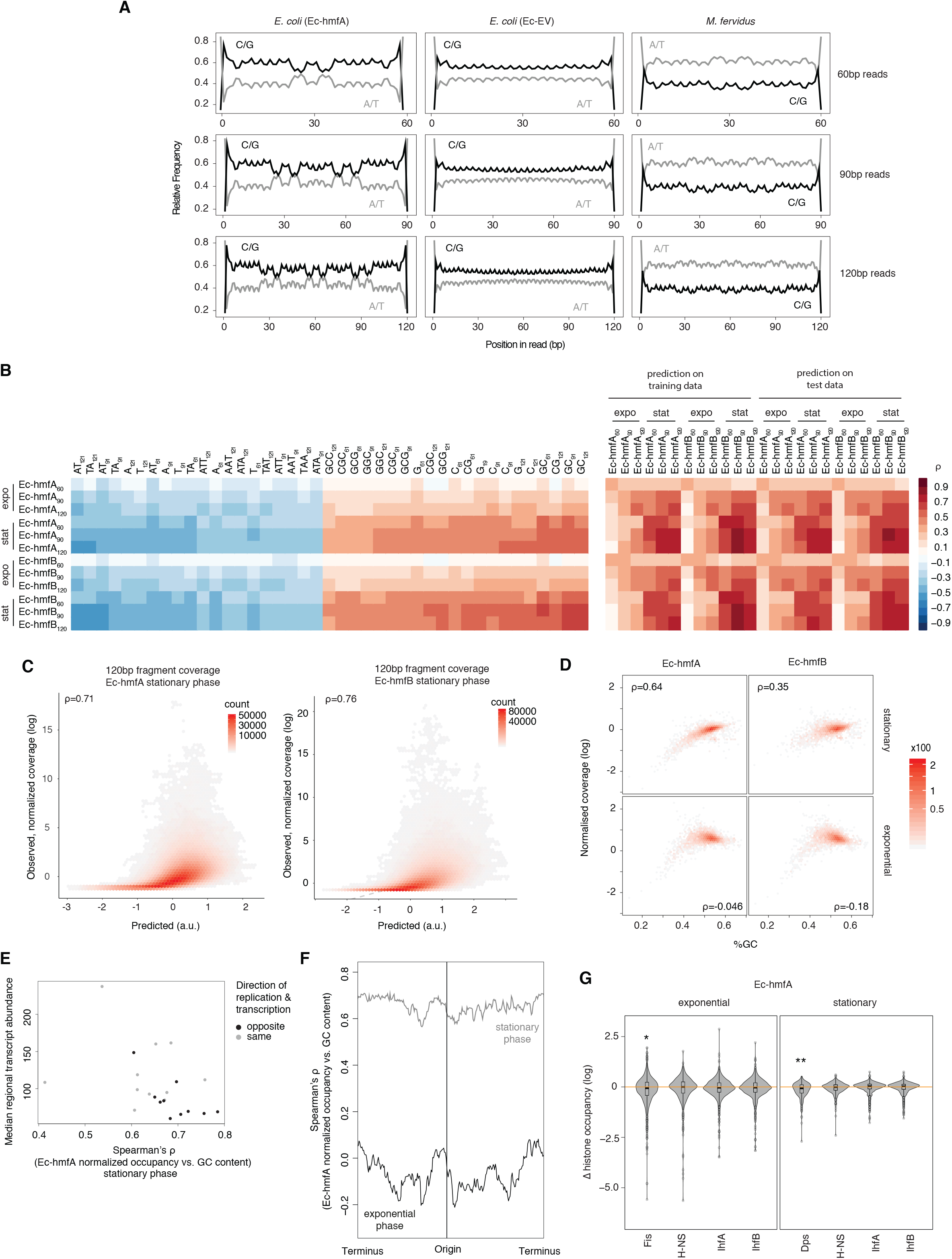
Sequence and other predictors of histone occupancy in *E. coli*. **A.** Read-interal nucleotide enrichment profiles for reads of exact length 60/90/120bp.Symmetric enrichments are evident for Ec-hmfA and *M. fervidus* native fragments but not Ec-EV. **B.** Left panel: top and bottom 20 individually most informative k-mers to predict fragment size-specific normalized histone occupancy in different strains. Red and blue hues indicate positive and negative correlations between k-mer abundance and normalized occupancy, respectively. Right panel: performance of the full LASSO model on training and test data (see Methods). **C.** Correlations between predicted and observed coverage of 120±5bp fragments predicted at single-nucletoide resolution across the genome. All P<0.001. **D.** GC content and normalized coverage are positively correlated in stationary but not exponential phase. All P<0.001. **E.** The correlation between GC content and occupancy is stronger in genomic regions where transcriptional output is lower. Regional transcriptional output is computed as median transcript abundance in a 200-gene window. To assess potential interactions between replication and transcription, windows are computed separately for genes where the directions of transcription and replication coincide and those where they differ. **F.** The strength of the correlation between GC content and occupancy varies along the *E. coli* chromosome. Correlations are computed for 500 neighbouring genes using a 20-gene moving window. **G.** Histone occupancy in regions previously found to be bound or unbound by a particular nucleoid-associated protein in *E. coli*. Δ histone occupancy is defined as the difference in histone occupancy in a region bound by a given NAP and the nearest unbound region downstream. Negative Δ histone occupancy values therefore indicate greater histone occupancy in areas not bound by the focal NAP, suggestive of competition for binding or divergent binding preferences. *P<0.005 **P<0.001

We suspect that stationary phase represents a comparatively more settled state, characterized by reduced replication, transcription, and other DNA-templated activity, that is more conducive to the establishment or survival of larger oligomers (Figure 1B) and where nucleosome formation is better able to track intrinsic sequence preferences. In support of this hypothesis, we find that transcriptional activity modulates the relationship between GC content and occupancy: the relationship is stronger where transcriptional activity is weaker (ρ=−0.46, P=0.039; Figure 3E). We also find that the strength of the correlation varies along the genome in a fashion suggestive of replication-associated biases, being relatively stronger, on average, further away from the origin of replication, particularly in exponential phase (Figure 3F). In contrast, local competition with endogenous nucleoid-associated proteins (NAPs) appears to have a minor impact on histone binding patterns. Locations previously identified as bound by IhfA, IhfB, or H-NS (Prieto *et al*, 2012; Kahramanoglou *et al*, 2011) are occupied by nucleosomes to the same extent as neighbouring unbound regions, indicating that histones are not significantly excluded from regions bound by these NAPs (Figure 3G). Histone occupancy in regions previously identified as bound by Dps, a NAP exclusively but abundantly expressed during stationary phase, is lower than in neighbouring unbound regions, consistent with a model where Dps and histones compete for some binding sites (Figure 3G). Effect size, however, is small. The same is true for Fis, which is expressed during the early stages of the growth cycle. In summary, we find that HMfA and HMfB readily form nucleosomes on the *E. coli* genome and do so in a manner consistent with sequence as a key determinant of positioning and occupancy, modulated by transcription and replication.

### Evidence that nucleosome formation locally represses transcription

Next, we asked whether histone occupancy in *E. coli* affects transcription. In particular, we wanted to establish whether histone binding to DNA exerts global or local repressive effects. To address these questions, we generated two additional strains, Ec-hmfA_nb_ and Ec-hmfB_nb_, where *hmfA* and *hmfB*, respectively, were recoded to carry three amino acid changes (K13T-R19S-T54K) previously shown to abolish DNA binding of HMfB (Soares *et al*, 2000). MNase treatment of these strains resulted in digestion profiles similar to Ec-EV, consistent with compromised DNA binding (Figure S3). Using RNA-Seq, we quantified differential transcript abundance in Ec-hmfA versus Ec-EV and Ec-hmfA_nb_ versus Ec-EV (see Methods) and then excluded genes from further analysis that were significantly up-regulated (or down-regulated) in both comparisons, reasoning that coincident patterns of change likely derive from systemic responses to heterologous expression and are not uniquely attributable to binding. We then considered differential expression in Ec-hmfA/B versus Ec-EV for the remaining genes as a function of nucleosome occupancy.

Looking at normalized coverage across gene bodies, annotated promoters and experimentally mapped transcriptional start sites, we find evidence for nucleosome-mediated dampening of transcriptional output. Notably, genes that are significantly (P_adj_<0.05) down-regulated in histone-bearing strains display significantly higher nucleosome occupancy at TSSs than upregulated genes (Figure 4A, Wilcoxon test). This is true regardless of whether we consider occupancy at a single base assigned as the TSS, occupancy in a ±25bp window around that site, or occupancy across annotated promoters (see Methods). This signal is lost almost entirely when considering a promoter-proximal 51bp control window centred on the start codon (Figure S4), suggesting that the effect is locally specific. The relationship between transcriptional changes and average histone occupancy across the gene body is more complex as weaker effects in the expected direction are evident for Ec-hmfA but not Ec-hmfB (Figure S4). Interestingly, repressive effects at TSSs in particular appear to be driven by larger oligomeric nucleosomes (90bp, 120bp, 150bp, Figure 4B, Figure S4), perhaps suggesting that larger oligomers are more significant barriers to transcription initiation and elongation. This might be so either because larger oligomeric complexes are intrinsically more stable and hence harder to bypass/displace or because, akin to H-NS, larger oligomers can extend across and occlude the promoter from an initial point of stable nucleation, reaching into sequence territories that are rarely populated independently by smaller complexes (Henneman *et al*, 2018; Hocher *et al*, 2019).

**Figure 4.**
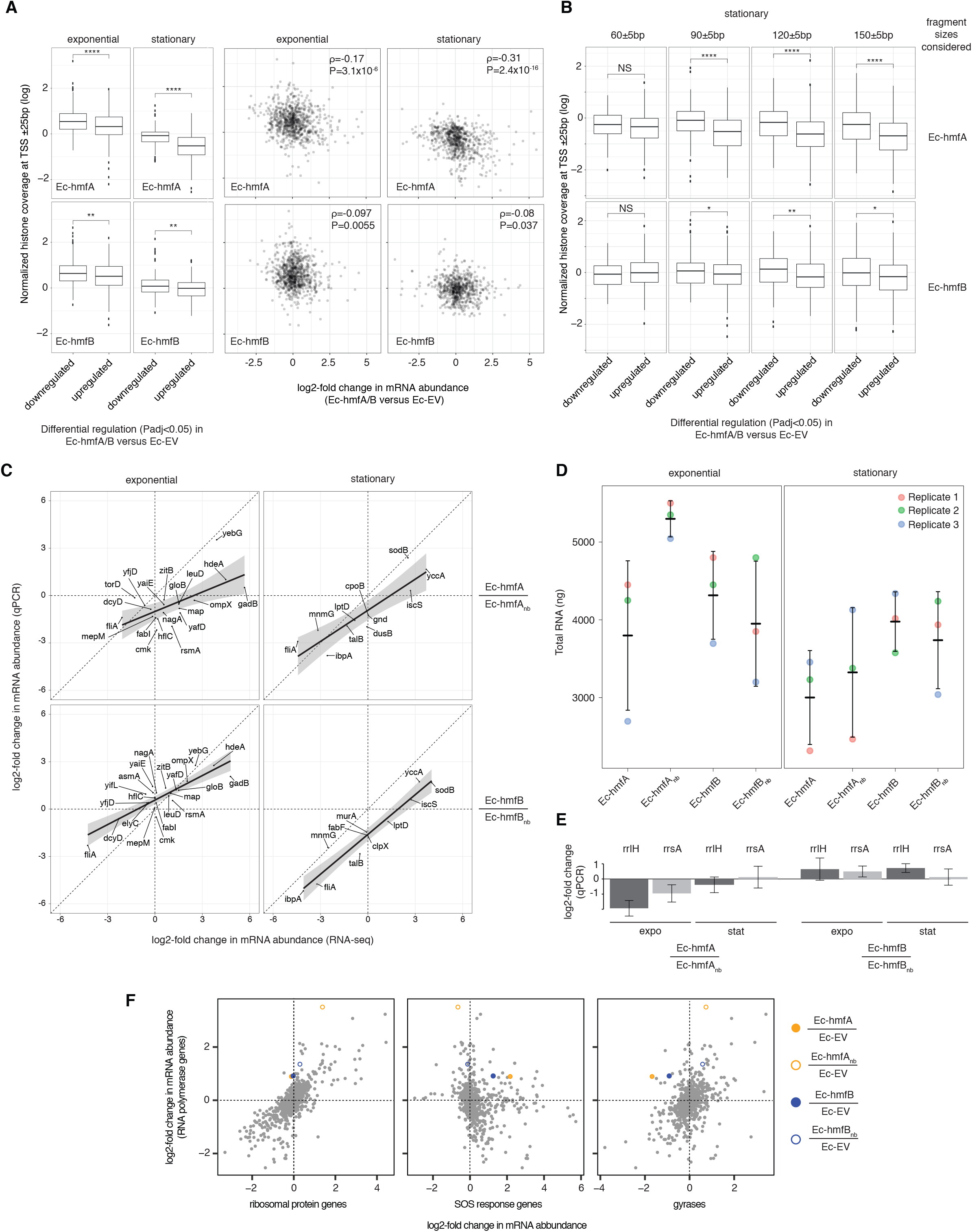
The impact of archaeal histones on transcription in *E. coli*. **A.** Reduced transcript abundance in histone-expressing strains is associated with higher average histone occupancy at the TSS.Top panels: Ec-hmfA. Bottom panels: Ec-hmfB **B.** Genes that are significantly downregulated in histone-expressing strains exhibit higher coverage of large (90+bp) but not small (60bp) fragments.Top panels: Ec-hmfA. Bottom panels: Ec-hmfB **C.** Relative changes in transcript abundance comparing histone-expressing and non-binding-histone-expressing strains, as measured using RNA-Seq and qPCR. As yeast RNA was used for spike-in normalization, shifts away from the diagonal can be interpreted as differences in RNA abundance between strains. For example, for stationary Ec-hmfB/Ec-hmfB_nb_, all but one gene measured show lower abundance in Ec-hmfB compared to Ec-hmfB_nb_. **D.** Total RNA quantification for binding and non-binding strains. **E.** Relative changes in the abundance of two ribosomal rRNA genes as measured by qPCR. **F.** Differential expression in histone-expressing strains compared to the empty vector control in the context of differential expression responses observed in previous RNA-Seq experiments (see Methods). Ec-hmfA_nb_ exhibits extreme upregulation of RNA polymerase genes, while SOS genes are upregulated and gyrases downregulated in strains with DNA-binding histones but not in strains carrying mutant histones. ****P<0.001; ***P<0.005; **P<0.01; *P<0.05; expo: exponential phase; stat: stationary phase.

### Do archaeal histones dampen transcription globally?

While the quantitative relationship between nucleosome occupancy and transcriptional change at individual loci is comparatively weak (Figure 4A), it is conceivable that histones might dampen transcriptional output globally and proportionately. Such blanket repression would not necessarily be evident from our RNA-Seq data, which did not include spike-ins, and we might therefore underestimate histone-mediated repression. To explore this possibility, we quantified total RNA abundance for a defined number of cells and carried out qPCR experiments for a small collection of mRNAs using yeast RNA for spike-in normalization (see Methods). Reassuringly, fold-changes inferred from qPCR and RNA-Seq show high levels of correspondence (Figure 4C). More interestingly, there appears to be a tendency for lower mRNA abundance in binding versus non-binding strains under at least some conditions (Ec-hmfB during stationary phase and Ec-hmfA during exponential phase). Taken at face value, this observation is consistent with a model where histones exert a global dampening effect on gene expression. However, things are more complicated than they appear. Total RNA in Ec-hmfA_nb_ (and rRNA abundance as its likely principal driver) are strongly and unexpectedly elevated compared to all other strains (Figure 4C-D). Rather than global nucleosome-mediated repression in Ec-hmfA, this is more compatible with global upregulation in Ec-hmfA_nb_. To investigate this further, we compared our RNA-Seq data to a large collection of *E. coli* K-12 differential expression profiles covering a variety of environmental and genetic perturbations (see Methods). Considering, in this broader context, the fold changes observed in our binding and non-binding strains versus Ec-EV, we find that genes involved in the SOS response are upregulated whereas expression of gyrases is downregulated in Ec-hmfA and Ec-hmfB but not in Ec-hmfA_nb_ and Ec-hmfB_nb_, highlighting system-wide (stress) responses specific to histone binding (Figure 4F).

Most notably, however, Ec-hmfA_nb_ stands out with regard to the expression of RNA polymerase genes. Normally, there is a strong relationship, across conditions and perturbations, between ribosomal and RNA polymerase genes (Pearson’s r=0.76, P<2.2×10^−16^, Figure 4F), reflecting concerted regulation in the context of growth (up) and stress (down). RNA polymerase levels in Ec-hmfA_nb_ versus Ec-EV during exponential phase, however, are disproportionately high (Figure 4F). While we do not currently know the reason for excess RNA polymerase expression, this observation is important because it highlights a more general point: global up- or downregulation need not reflect distributed effects in *cis* – such as concurrent downregulation of transcription at multiple loci because of genome-wide nucleosome formation – but might instead be caused in *trans* by altered expression of regulatory factors with global reach such as RNA polymerases. Importantly, however, we note that the unusual behaviour of Ec-hmfA_nb_ does not challenge the evidence for local repressive effects reported above (Figure 4A-B), as the correlations between nucleosome occupancy at promoters and transcript abundance were computed by comparing Ec-hmfA versus Ec-EV.

### Histone binding is associated with mild morphological and growth defects

Despite widespread transcriptional changes, gross cell morphology and growth rate appear surprisingly normal. Histone-expressing cells are longer than Ec-EV cells, particularly in stationary phase, but appear to divide normally (Figure 5A-B). They also grow at remarkably similar rates to control strains that carry non-binding histones (Figure 5C), exhibiting only transient reductions in growth rate following induction, indicative of mild stress and consistent with elevated expression of SOS response genes (Figure 4F). Overall, growth of *E. coli* expressing DNA-binding archaeal histones is remarkably unremarkable.

**Figure 5.**
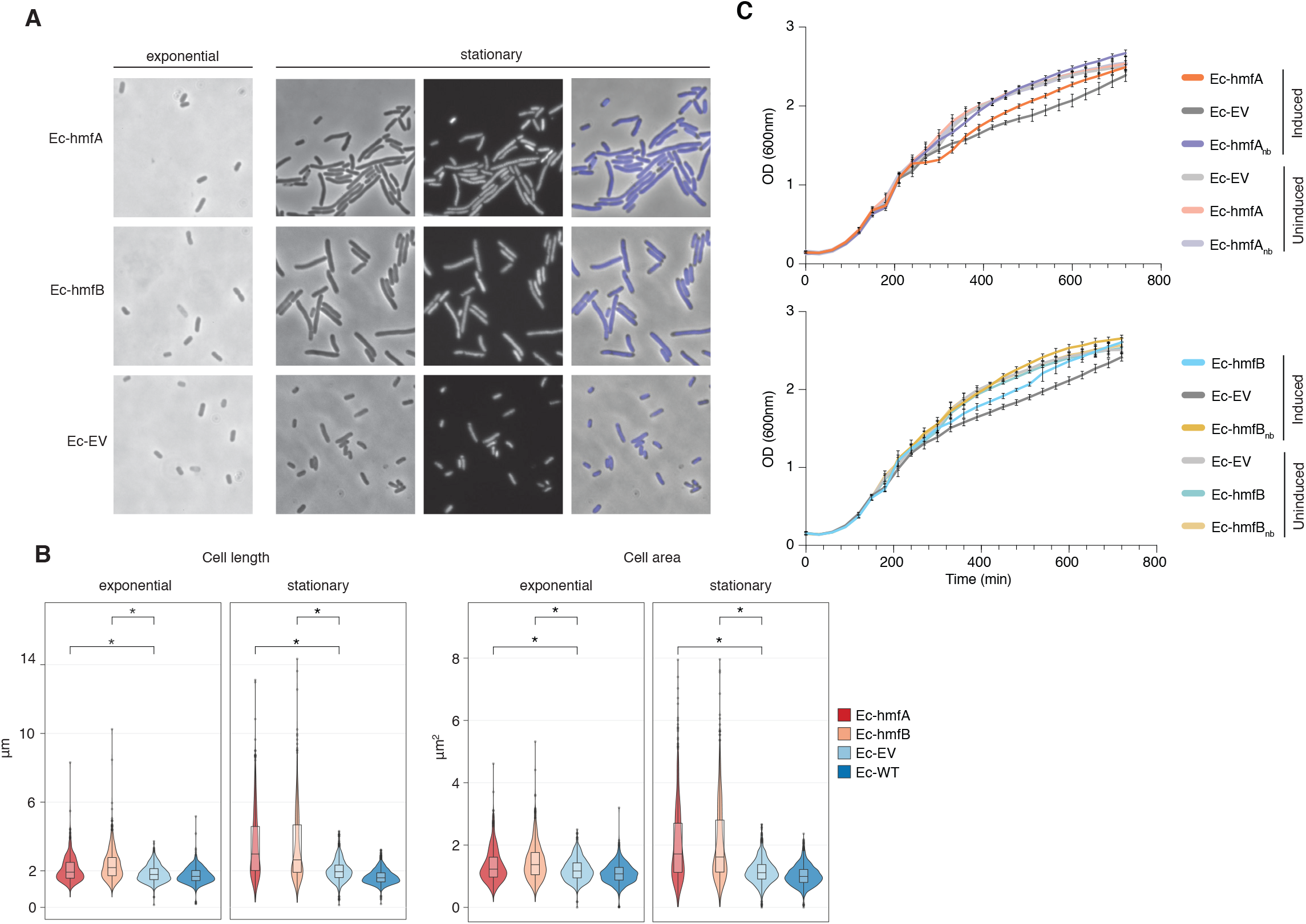
The impact of archaeal histones on *E. coli* growth. **A.** Morphological changes are triggered by HMfA and HMfB histones expression. Compared to the empty vector control, transformant Ec-hmfA and Ec-hmfB become significantly longer, particularly towards the final stage of cell cycle. DAPI staining suggests that the increase in cell length is not due to impaired cell division. Magnification 100x **B.** Quantification of cell length and area in histone-expressing and control strains. Some unexpectedly low values are likely attributable to debris being misidentified as cells. P<0.0001 **C.** Growth curves for induced and uninduced histone-expressing and control strains. Rhamnose was added for induction at 200min.

## DISCUSSION

The experiments reported above demonstrate that archaeal histones are surprisingly well tolerated when expressed in *E. coli*, a system that has not evolved to deal with nucleosomal structures. Despite binding ubiquitously to the *E. coli* genome, they do not fundamentally compromise critical DNA-templated processes. In particular, while we find some evidence that nucleosome occupancy locally restricts the output of the transcription machinery, gene expression is insufficiently perturbed to critically affect growth. In as much as histones will have non-linear, concentration-dependent effects on genome function, however, we note that increasing histone dosage might well force more severe repressive effects. The fact that, for practical reasons, we have explored only a small section of dosage space is therefore an obvious limitation of the current study.

In our experiments as well as during evolution, global wrapping of DNA into nucleosomes was likely facilitated by two factors in particular: First, by virtue of their AT-rich nature, promoters remain comparatively accessible to the transcription machinery, even in a naïve prokaryote whose sequence did not co-evolve to accommodate histones (Figure 2). Nucleosome-free regions at the TSS, a key features of nucleosome architecture in eukaryotes, might therefore have emerged, in the first instance, as a simple consequence of promoter composition. Once established, nucleosomes bordering the TSS were uniquely positioned to be co-opted into gene regulatory roles in eukaryotes and perhaps along different archaeal lineages, with nucleosome positioning later refined by evolution at specific loci to provide more nuanced control over transcriptional processes. Second, compared to their eukaryotic counterparts, archaeal nucleosomes appear to be more surmountable barriers to transcription elongation. Even at high histone concentrations, transcription through a HMf-chromatinized template *in vitro* is slowed but not aborted (Xie & Reeve, 2004), in line with the absence of recognizable histone remodelers from archaeal genomes. Thus, near-global coating of the genome with archaeal(-like) histone proteins might have evolved without severe repercussions for basic genome function before a more restrictive arrangement, perhaps coincident with the advent of octameric histone architecture, took hold during eukaryogenesis. From an evolutionary point of view, one might therefore call the ground state mediated by archaeal histones proto-restrictive.

To what extent restrictive, proto-restrictive, or permissive ground states exist in different archaea *in vivo* remains unclear. Experiments with histones from *M. fervidus*, *Methanococcus jannaschii*, and *Pyrococcus furiosus* have shown that archaeal nucleosomes can interfere with transcription initiation and elongation *in vitro* (Wilkinson *et al*, 2010; Soares *et al*, 1998; Xie & Reeve, 2004). However, significant inhibitory effects were only observed at high histone:DNA ratios (close to or above 1:1). Ratios of that magnitude, while regularly found in eukaryotes, need not be prevalent in archaea. Direct measurements of histone:DNA ratios are scarce and variable, with prior estimates in *M. fervidus* reporting stoichiometries as high as 1:1 (Pereira *et al*, 1997) and as low as 0.2-0.3:1 (Stroup & Reeve, 1992). Considering transcript levels as a (really rather imperfect) proxy, histones appear very abundant in *Thermococcus kodakarensis* and *Methanobrevibacter smithii* (Figure S5), strengthening the case for histones as global packaging agents in these species. In contrast, histone mRNAs are much less plentiful in *Haloferax volcanii* and *Halobacterium salinarum* (Figure S5), where less than 40% of the chromosome is resistant to MNase digestion (Takayanagi *et al*, 1992). Thus, histone:DNA stoichiometry likely varies substantially across taxa as well as along the growth cycle (Takayanagi *et al*, 1992; Dinger *et al*, 2000; Sandman *et al*, 1994).

Attempts to delete histone genes have also revealed considerable diversity across archaea. Histones are required for viability in *T. kodakarensis* and *Methanococcus voltae* (Cubonova *et al*, 2012; Heinicke *et al*, 2004), but can be removed with surprisingly muted effects on transcription in *Methanosarcina mazei* (Weidenbach *et al*, 2008) and *H. salinarum* (Dulmage *et al*, 2015). In both species, a comparatively small number of transcription units were affected by histone deletion, the majority of which was down- rather than upregulated. Taken together, these observations suggest that histones likely play a more variable, species-and context-dependent role in archaea, may only sometimes act as global repressive agents and, more generally, that care should be taken in projecting properties of eukaryotic histones onto those of archaea. In many instances, archaeal histones might be better understood with reference to bacterial NAPs, especially when considering how concentration drives opportunities for oligomerization, cooperativity, and bridging interactions with DNA. In this context, we note that our results are reminiscent of a recent study by Janissen and colleagues, who found that *dps* deletion in *E. coli* results in nucleoid decompaction but does not greatly impact transcription (Janissen *et al*, 2018). This provides some generality to the notion that architectural DNA-binding proteins, even if they bind to most of the genome and alter its compaction and gross structure, need not unduly interfere with transcription. The same study also highlights that, while polymerases may continue to access DNA and operate as usual, the same need not be true for other DNA-binding proteins: Dps substantially reduced the ability of several restriction enzymes to recognize and cut their target sites. Whether archaeal histones have similar effects in *E. coli* (beyond their ability to protect from MNase treatment), remains to be established. However, access regulation outside of a transcriptional context might well have provided the original evolutionary impetus for histones to spread across the genome, as genomes evolved to defend themselves against selfish elements that target the host genome for integration (Talbert *et al*, 2019). We note in this regard that our chromatinized *E. coli* strains might be of use for future synthetic biology applications. As more complex, combinatorial control of gene expression becomes a desirable genome engineering objective, limiting access to desired target sites will become an increasingly important design consideration (Cardinale & Arkin, 2012), as will chassis integrity in the face of potential invaders. As we find interference with transcription and replication to be limited, it will be interesting to experiment with expressing archaeal histones to restrict global access to the genome for specific DNA-binding factors or protect the genome against selfish element invasion (Sultana *et al*, 2019; Aslankoohi *et al*, 2012).

## METHODS

### Plasmid design

The coding sequences of *hmfA* and *hmfB* were codon-optimised for *E. coli* and synthesised as part of a rhamnose-inducible pD861 plasmid (Figure S1) by ATUM (Newark, CA). Originally, both plasmids also encoded a chromogenic protein to enable visual screening for induction. However, as the chromogenic protein was expressed at very high levels (Figure S2) and since we did not want to unduly increase cellular burden we removed the corresponding gene to yield pD861-hmfA. To generate non-binding histone mutants, *hmfA/hmfB* sequences were re-coded to carry three changes (K13T-R19S-T54K), previously shown to jointly abolish DNA binding of *hmfB* (Soares *et al*, 2000). These sequences were codon-optimized, synthesized and integrated onto a pD861 plasmid as above, without the chromogenic proteins, as was *hmfB*, for which cloning had failed. Plasmids pD861-hmfA, pD861-hmfB, pD861-hmfA_nb_, and pD861-hmfB_nb_ are identical expect for the sequences of the respective histone genes. *hmfA* was removed from pD681-hmfA to obtain Ec-EV.

### Bacterial transformation and growth

*E. coli* K-12 MG1655 cells were transformed via heat-shock with either pD861-hmfA, pD861-hmfB or pD861-EV, or the non-binding histone mutants pD861-hmfA_nb_ or pD861-hmfB_nb_ to generate strains Ec-EV, Ec-hmfA, Ec-hmfB, Ec-hmfA_nb_ and Ec-hmfB_nb_, respectively. All strains were grown in LB medium plus kanamycin (50μg/ml) at 37°C with agitation (170 rpm). Histone expression was induced by adding L-Rhamnose monohydrate to a final concentration of 15mM at OD600 ~0.6. Cells were harvested after 2hrs or 16-17hrs following induction.

### Protein purification

HMf protein purification was performed as in (Starich *et al*, 1996).

### Coomassie staining

Bacteria were harvested by centrifugation (4000rpm for 15min at 4°C), the supernatant discarded, and the pellet resuspended in a small volume of Histone Wash Buffer (50mM Tris-HCl pH 7.5, 100mM NaCl, 1mM EDTA). Cell envelopes were disrupted using a Bioruptor Plus sonication system (Diagenode s.a., Belgium) for 10 cycles, 30 seconds on/off with power set to high. The soluble protein fraction was separated from cellular debris by centrifugation at 15000 x g for 15min at 4°C, while the insoluble fraction was obtained by re-suspending the pelleted debris in Histone Wash Buffer. The protein concentration in the cell lysate was quantified with a Pierce BCA Protein Assay Kit (ThermoFisher Scientific, UK) using the provided albumin as standard. Protein fractions were separated by means of 16.5% Tris-tricine precast gels (Bio-Rad Laboratories, California) and bands were revealed by colloidal Coomassie (InstantBlue, Sigma-Aldrich) staining. Histone-expressing strains showed a band close to the size expected for HmfA/B (Figure S2). This band was excised and protein identity confirmed as HMfA/B via mass spectrometry.

### Growth assays

Overnight pre-cultures were diluted 1:500 into LB medium plus kanamycin (50μg/ml). Samples were plated in replicate into a flat bottom Nunc 96-well plate (ThermoFisher Scientific, UK) and incubated at 37°C at 100rpm for 30min. OD measurement were performed using a high-throughput microplate reader (FLUOstar Omega, BMG LABTECH GmbH, Ortenberg, Germany) in which bacteria were grown at 37°C under continuous shaking (~500rpm, double orbital). Optical density was measured at 600nm every 30min for 12.5hrs. For induction, the microplate reader was paused at cycle 6 and L-Rhamnose monohydrate added manually to the relevant wells to a final concentration of 15mM. Results presented are from three biological replicates per strain, each averaged across six technical replicates.

### MNase digestion – *E. coli*

Bacterial cultures were harvested by centrifugation (4000rpm for 15min at 4°C), the supernatant discarded and the pelleted cells re-suspended in chilled 1x PBS (Gibco, ThermoFisher Scientific, UK). Cells were then fixed by adding a fixation solution (100mM NaCl, 50mMTris-HCl pH 8.0, 10% formaldehyde) for 10min at room temperature under slow rotation, after which fixation was quenched by adding 140mM glycine. Following a further round of centrifugation (4000rpm for 5min at 4°C), bacteria were washed twice with 10ml chilled 1x PBS and incubated in a lysozyme buffer (120mM Tris-HCl pH 8.0, 50mM EDTA, 4mg/ml Lysozyme) for 10min at 37°C to generate protoplasts. Cells were pelleted (15000rpm for 3min at room temperature) and re-suspended in 500μl of lysis buffer (10mM NaCl, 10mM Tris-HCl pH 7.4, 3mM MgCl2, 0.5% NP-40, 1x Pi, 0.15mM Spermine, 0.5mM Spermidine), transferred to a new microcentrifuge tube and incubated on ice for 20min. Subsequently, the lysate was spun down and the pellet washed with 500μl of -CA buffer (15mM NaCl, 10mM Tris-HCl pH 7.4, 60mM KCl, 1x Pi, 0.15mM Spermine, 0.5mM Spermidine) without re-suspending. The washed pellet was finally re-suspended in 500μl of +CA buffer (15mM NaCl, 10mM Tris-HCl pH 7.4, 60mM KCl, 1mM CaCl2, 0.15mM Spermine, 0.5mM Spermidine) to a uniform suspension. 50μl of this suspension were digested with micrococcal nuclease (LS004798, Worthington Biochemical Corporation, NJ; 500U/ml for Ec-hmfA and Ec-hmfB, 50U/ml for Ec-EV) for 10min (20 minutes for cells in stationary phase) at room temperature and finally blocked with a STOP solution containing calcium-chelating agents (100mM EDTA, 10mM EGTA). Each sample was further diluted with -CA buffer and treated with 10% SDS and 150ng/ml proteinase K overnight at 65°C with shaking at 500rpm. Undigested DNA fragments was purified by two rounds of phenol:chloroform extraction separated by an RNase A digestion step (100μg/ml, 2h at 37°C with shaking at 500rpm). Finally, DNA fragments were precipitated in ethanol and re-suspended in 40μl distilled water. The quality of the digest and the size of the retrieved fragments were assessed by agarose DNA electrophoresis (2.5% agarose gel in 1x TBE run at 150V for 30min).

### MNase digestion – *M. fervidus*

Frozen pellets of *M. fervidus* harvested in late exponential and stationary phase were purchased from the Archaeenzentrum in Regensburg, Germany. We then followed the MNase protocol outlined above with the following modifications: first, ~0.5g of pellet were thawed and re-suspended in 9ml of 1x PBS before fixation. Second, due to differences in cell wall composition between *M. fervidus* and *E. coli*, the lysozyme digestion step was replaced by mechanical disruption with a French press: after the wash that follows fixation, the pellet was re-suspended in 20ml of chilled 1x PBS, the cell suspension passaged twice through a TS Series French press (Constant Systems) at 15kpsi and then spun down at 4000rpm for 15 minutes at 4°C before proceeding with cell lysis. Finally, the extracted chromatin was re-suspended in 250μl of +CA buffer (instead of 500μl). Digestion, fragment purification, sequencing and analysis were performed as for *E.coli* but with a micrococcal nuclease concentration of 100 U/ml.

### MNase digest sequencing

Size distributions of the DNA fragments retrieved by MNase digestion of strains Ec-EV, Ec-hmfA, Ec-hmfB and *M. fervidus* were analysed with an Agilent Bioanalyser DNA1000 chip. For each of these strains, three biological replicates were selected for sequencing. Twenty nanograms per sample were used for library construction with the NEBNext Ultra II DNA Library Prep Kit for Illumina and NEBNext Multiplex Oligos for Illumina. The output was then taken to 10 PCR cycles and purified using a 1.8x Ampure XP bead clean-up kit. Libraries were quantified via Qubit and quality assessment carried out on an Agilent Bioanalyser DNA 1000 chip. Libraries were then sequenced on an Illumina MiSeq sequencer using single-end 160bp reads.

### Read processing

Reads were trimmed using Trimmomatic-0.35 (single-end mode, ILLUMINACLIP:2:30:10) to remove adapter sequences. This did not remove short remnant adapter sequences so that we submitted reads to a further round of trimming using Trimgalore v0.4.1 with default parameters. Trimmed reads were aligned, as appropriate, to either the *Escherichia coli* K12 MG1655 genome (NC_000913.3) or the *M. fervidus* DSM2088 genome (NC_014658.1) using Bowtie2 (Ben Langmead & Salzberg, 2012). Only uniquely mapping reads were retained for further analysis. Per-base coverage statistics were computed using the genomeCoverageBed function bedtools2 suite (Quinlan & Hall, 2010). For certain downstream analyses, aligned reads were separated into bins of different length, each corresponding to DNA fragments wrapped around histone oligomers of increasing lengths (60±5bp, 90±5bp, etc.).

### Peak calling

Nucleosome peaks were called using the NucleR package in R as described previously (Hocher *et al*, 2019). See Table S2 for the relevant Fourier parameters.

### LASSO modelling

LASSO modelling was carried out for different footprint size classes (60±5bp, 90±5bp, 120±5bp) using empty vector-normalized coverage. Empty vector coverage was computed across fragment sizes and coverage across the genome uniformly increased by 1 to enable analysis of zero-coverage regions. K-mer counts (k={1,2,3,4}) were computed using the R seqTools package over windows of 3 different sizes (61bp, 91bp, 121bp). Subsequent LASSO modelling was then carried out as described previously (Hocher *et al*, 2019), with models trained on one sixth of the *E.coli* genome (genomic positions 0-773608) and tested on the remainder of the genome.

### Transcriptional start sites

Experimentally defined transcriptional start sites were obtained from RegulonDB (Salgado *et al*, 2013) (http://regulondb.ccg.unam.mx/menu/download/datasets/files/High_throughput_transcription_initiation_mapping_with_5_tri_or_monophosphate_enrichment_v3.0.txt). The position inside each broad TSS associated with the most reads (column 3 in the file above) was defined as the TSS for downstream analysis. Promoter annotations were obtained from the same source (http://regulondb.ccg.unam.mx/menu/download/datasets/files/PromoterSet.txt).

### Comparison with other transcriptomes

All available transcriptomic data corresponding to *E.coli* K-12 strains were downloaded from the *E. coli* Gene Expression Database (GenExpdb, https://genexpdb.okstate.edu), which aggregates differential transcriptional responses (increased/decreased mRNA expression computed from pairwise comparisons in different individual studies). To simplify analysis, we computed a single mean fold-change value across genes in certain categories of interest (ribosomal, RNA polymerase, and SOS response genes). Specifically, these were *rpoA, rpoB* and *rpoC* (RNA polymerase category), all *rps*\* and *rpl*\* genes (ribosomal genes) and *dinB*, *dinD*, *sulA*, *recA*, *sbmC*, *recN*, previously defined as SOS-inducible genes (Khil & Otero, 2002).

### RNA extraction and sequencing

250μl of culture were harvested from late exponential and stationary phase by centrifugation (15000 x g at 4°C for 15min). The supernatant was discarded and the pellet re-suspended in 100μl of Y1 Buffer (1M Sorbitol, 0.1M EDTA, 1mg/ml lysozyme, 0.1% β-mercaptoethanol) and incubated at 37°C for 1h at 500rpm. The cell suspension was added to 350μl of RLT buffer, 250μl 100% ethanol and loaded onto an RNeasy column from the RNeasy Kit (Qiagen, Germany). RNA was then washed and eluted following the manufacturer’s protocol. Eluted samples were incubated with DNase I (New England Biolabs, MA) for 10min at 37°C and then cleaned up with a second passage through the RNeasy column (loading, washes and elution according to manufacturer’s instructions). Samples were finally eluted in 30μl of RNase-free water and RNA quantified with Nanodrop. Quality assessment of the extracted RNA was carried out with an Agilent Bioanalyser RNAnano chip and five replicates per strain/condition were chosen for sequencing.

### RNA sequencing

For each replicate/strain/condition, 1.5μg of total RNA were depleted of rRNA using the Ribo-Zero rRNA depletion kit (Illumina) and libraries constructed using a TrueSeq Stranded RNA LT Kit (Illumina). After 12 PCR cycles, library quality was assessed with an Agilent Bioanalyser HS-DNA chip and quantified by Qubit. No size selection was carried out and the samples were sequenced on a HiSeq 2500 machine using paired-end 100bp reads.

### Transcriptome analysis

Using Bowtie2, reads were first aligned to all annotated non-coding RNA genes (rRNA, tRNA, etc.). Reads that mapped to any of these genes were discarded, even if they mapped to more than one location in the genome. We then used Trim Galore v0.4.1 with default parameters to trim adapters and low quality terminal sequences. Trimmed reads were then aligned to the *E. coli* K12 MG1655 genome (NC_000913.3) with Bowtie2 (--no-discordant --no-mixed). As a technical aside, we note that, despite the above filtering steps, some of the samples had an unusually low alignment rate (<30%). We found that most of the unaligned reads were perfect matches to rRNA sequences from *Bacillus subtilis* but not *E. coli* and had therefore eluded the above filter. As contamination at this scale is unlikely (no bacteria other than *E. coli* are grown or sequenced in the lab and a plain LB control was added to check for contamination when growing the samples for RNA extraction), we suspect the these reads are the result of carrying over RiboZero oligos. The addition of a further round of filtering to discard reads that match non-coding RNA sequences from *Bacillus subtilis* increased the alignment rate to *E. coli* index up to ~90%.

By-gene read counts were computed from read alignments using the summarizeOverlaps function (mode=“Union”, singleEnd=FALSE, ignore.strand=FALSE, fragments=TRUE) from the GenomicAlignments package in BioConductor. Differential gene expression analysis was carried out using DESeq2 (Love *et al*, 2014). Replicates found to be outliers in principal component analysis and that were subsequently excluded from differential expression analysis are listed in Table S3.

### Total RNA quantification

Three biological replicates for strains Ec-hmfA, Ec-hmfB, Ec-hmfA_nb_ and Ec-hmfB_nb_ were harvested for total RNA extraction. Specifically, ~2×10^8^ cells/replicate in late exponential and ~4×10^8^ cells/replicate in stationary phase were incubated with RNAprotect Bacteria Reagent (Protocol 3, Qiagen, Germany) according to manufacturer’s instructions. Cells were pelleted by centrifugation at 5000 x g at 4°C for 10min, the supernatant was decanted and the pellets snap frozen in liquid nitrogen and stored at −80°C. Single colonies of *S. cerevisiae* strain BY4741, a gift from Luis Aragón Alcaide, were used to inoculate 5ml overnight cultures. Fresh cultures were inoculated at OD ~0.2 and grown until OD ~1. Pellets corresponding to 0.5 OD of cells were snap frozen in liquid nitrogen. After thawing, one yeast aliquot was added to each bacterial pellet in a final volume of 700μl of RLT buffer (RNeasy mini Kit). The re-suspended cellular mixture was transferred to a tube containing acid washed glass beads (equal amounts of 425-600μm and 710-1180μm beads) and mechanically disrupted using a TissueLyser machine (Qiagen, Germany) with two successive cycles of 5min and 3min at maximum speed. Tubes were spun at 15000rpm for 30s to settle re-suspended beads and 500μl of the obtained lysate were transferred into a new tube containing 500μl of 100% ethanol. RNA purification was carried out using an RNeasy Kit (Qiagen, Germany) following manufacturer’s instructions, including on-column DNase I digestion. DNase treatment of stationary samples was extended to 30min to reduce residual contamination from genomic DNA given the higher number of cells in the input. Samples were eluted in 30μl of RNase-free water and RNA quantified with Nanodrop. 850ng of total RNA/sample were retrotranscribed with Superscript III Reverse Transcriptase (Invitrogen) and the cDNA was subsequently diluted fifty fold.

### Quantitative Real-Time PCR

Primers (Table S4) were designed with IDT PrimerQuest. Per reaction, 5μl of cDNA were used and amplification of selected target genes was detected using PowerUp SYBR Green Master Mix (Applied Biosystems) according to manufacturer’s instructions, processing three technical replicates per sample. Yeast mRNAs were used as internal calibrator in each reaction (see primer table). qPCR was performed on 384-well plates in a QuantStudio 7 Flex system (Applied Biosystems) using the default “fast” cycling conditions and a total reaction volume of 12μl. Primer specificity was evaluated based on qPCR product melting curve analysis. Ct values were automatically calculated by QuantStudio Real-Time PCR Software v1.2.

### Microscopy

Overnight pre-cultures of Ec-EV, Ec-hmfA, and Ec-hmfB were diluted in fresh LB medium plus antibiotic and grown as described above. ~300μl of culture were harvested by centrifugation (15000rpm for 15min). Pellets were resuspended in 1% FA in PBS and fixed for 10min at room temperature. Fixating agent was removed by spinning (15000rpm for 15 min) and pellets were resuspended in 1ml PBS. 5μl of cellular suspension was spread onto an agarose pad, covered in VectaShield containing DAPI (Vector Laboratories) and the excess liquid removed. Slides were imaged using a Manual Leica DMRB with phase contrast and DIC for transmitted light illumination. For quantification, images from three independent experiments were analysed with MicrobeJ (Ducret *et al*, 2016) to perform automatic cell detection and size measurements. MicrobeJ image profiles were manually curated to remove background and wrongly detected debris. For each sample/condition, measurements of cell length and area are derived from averages across ~10 independent pictures.

### NAPs binding regions

Genomic regions bound by Fis and H-NS were obtained from (Kahramanoglou *et al*, 2011), regions bound by IHF from (Prieto *et al*, 2012), and regions bound by Dps from (Antipov *et al*, 2017). Differential histone occupancy was computed between regions bound by a given NAP and the unbound region immediately downstream.

### Data availability

Datasets generated for this study have been deposited in the NCBI Gene Expression Omnibus under accession number GSE127680 (https://www.ncbi.nlm.nih.gov/geo/query/acc.cgi?acc=GSE127680).

## ACKNOWLEDGEMENTS

We would like to thank Ziwei Liang, Till Bartke, Ben Foster, and Kathleen Sandman for experimental advice, training, and sharing protocols; Finn Werner for his continued support and mentorship; Madan Babu, Ben Lehner, Peter Sarkies and members of the LMS Quantitative Biology section for discussions; Jacob Swadling for help with structure visualizations, and the MRC LMS Genomics and Proteomics facilities for sequencing and mass spectrometry. This work was supported by Medical Research Council core funding to TW.

## AUTHOR CONTRIBUTIONS

MR carried out all experiments and analyses. AH supported the analysis of MNase data, implemented comparative transcriptomic analysis, and carried out qPCR experiments alongside MR. MM provided training and co-supervised the project. TW conceived the study, supervised the project, participated in analysis and data interpretation and wrote the paper with input from all authors.

