## Supplementary Material for "Chromatinization of *Escherichia coli* with archaeal histones"

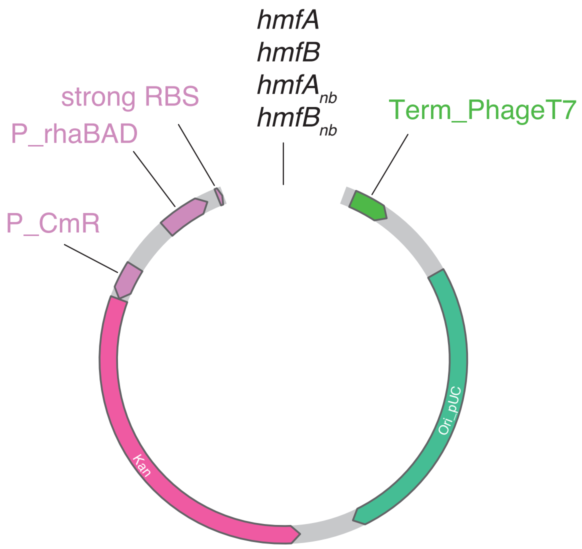

**Figure S1. Layout of pD681-derived plasmids used in this study**

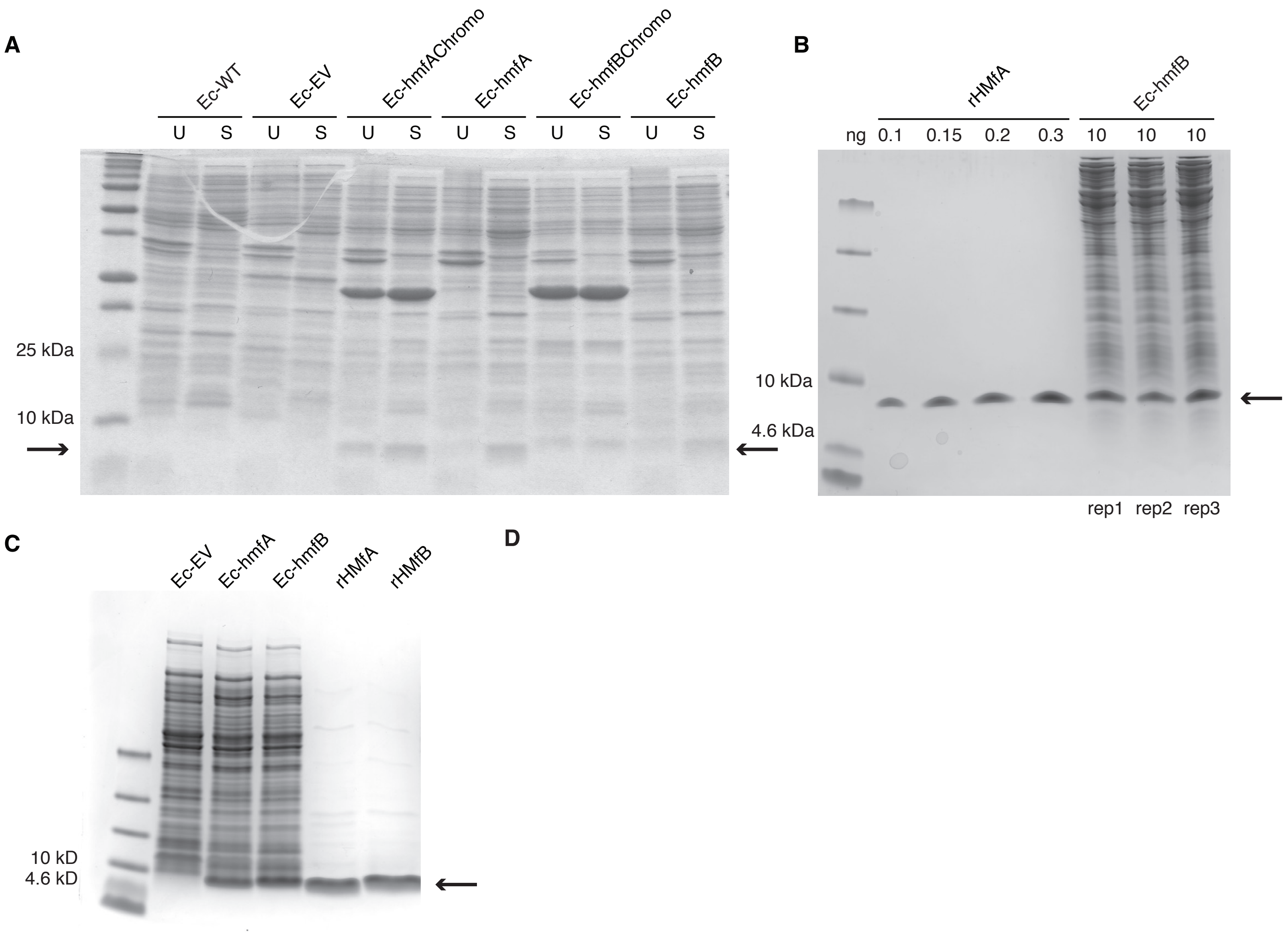

**Figure S2. Detection and quantification of HMf expression in *E. coli*. A.** Polyacrilamide gel comparing soluble (S) and unsoluble (U) fractions obtained by lysis of transformant strains (Ec-EV, Ec-hmfA, Ec-hmfB) and wild type *E. coli* (Ec-WT). Ec-hmfAChromo and Ec-hmfBChromo are histone-expressing strains that also bear a chromogenic protein. These were not further analyzed in this study (see Methods). **B.** HMfA/B protein levels in the transformant strain was estimated from SDS-PAGE bands by densiometry. 10ng of cell lysate was loaded onto a precast tris-tricine gel and the intesity of the band corresponding to HMfB (arrow) was compared to a standard curve made with increasing amounts of purified recombinant rHMfA (see Methods). **C.** Tris-tricine gel showing HMfA and HMfB expression in the soluble fraction of cell lysate and after Heparin-column purification. Ec-EV is included as negative control.

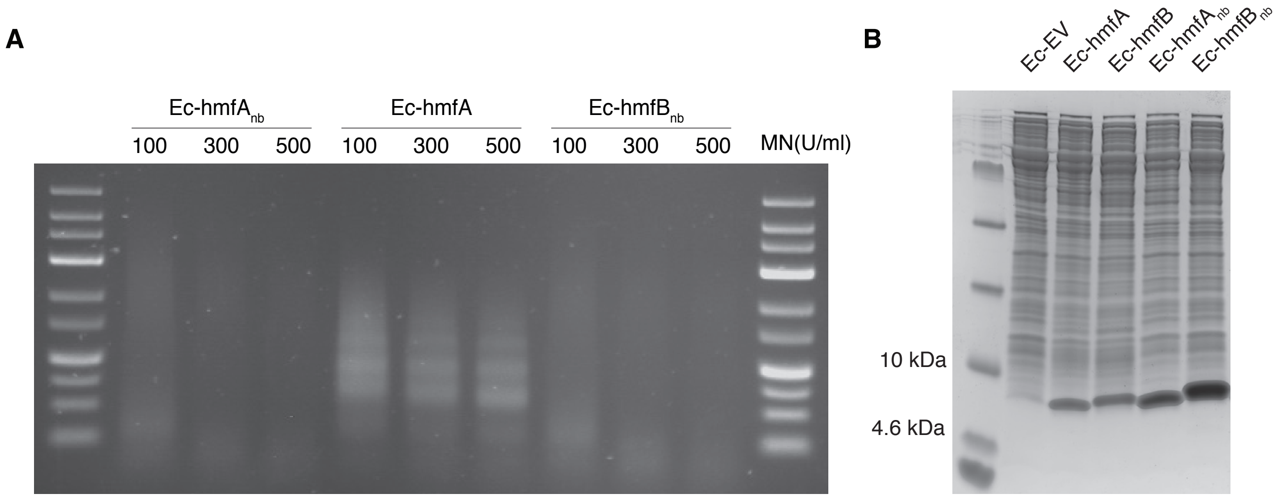

**Figure S3. Expression of non-binding histone mutants. A.** Agarose gel comparing the MNase digestion profiles of Ec-hmfA to strains expressing non-binding HMfA and HMfB mutants **B.** Coomassie staining of the soluble fractions of cell lysates from all transformant strains discussed in this study. Note that non-binding histone variants are - for unknown reasons - more highly expressed than their DNA-binding progenitors and might, by virtue of this higher expression and their different sequence, trigger qualitatively and quantitatively different responses. This makes an indirect comparison (comparing Ec-hmfA/B to Ec-EV after factoring out shared responses with Ec-hmf$A_{nb}$/$B_{nb}$ versus Ec-EV, as described in the main text) preferable to directly comparing differential expression in Ec-hmfA/B versus Ec-hmf$A_{nb}$/$B_{nb}$.

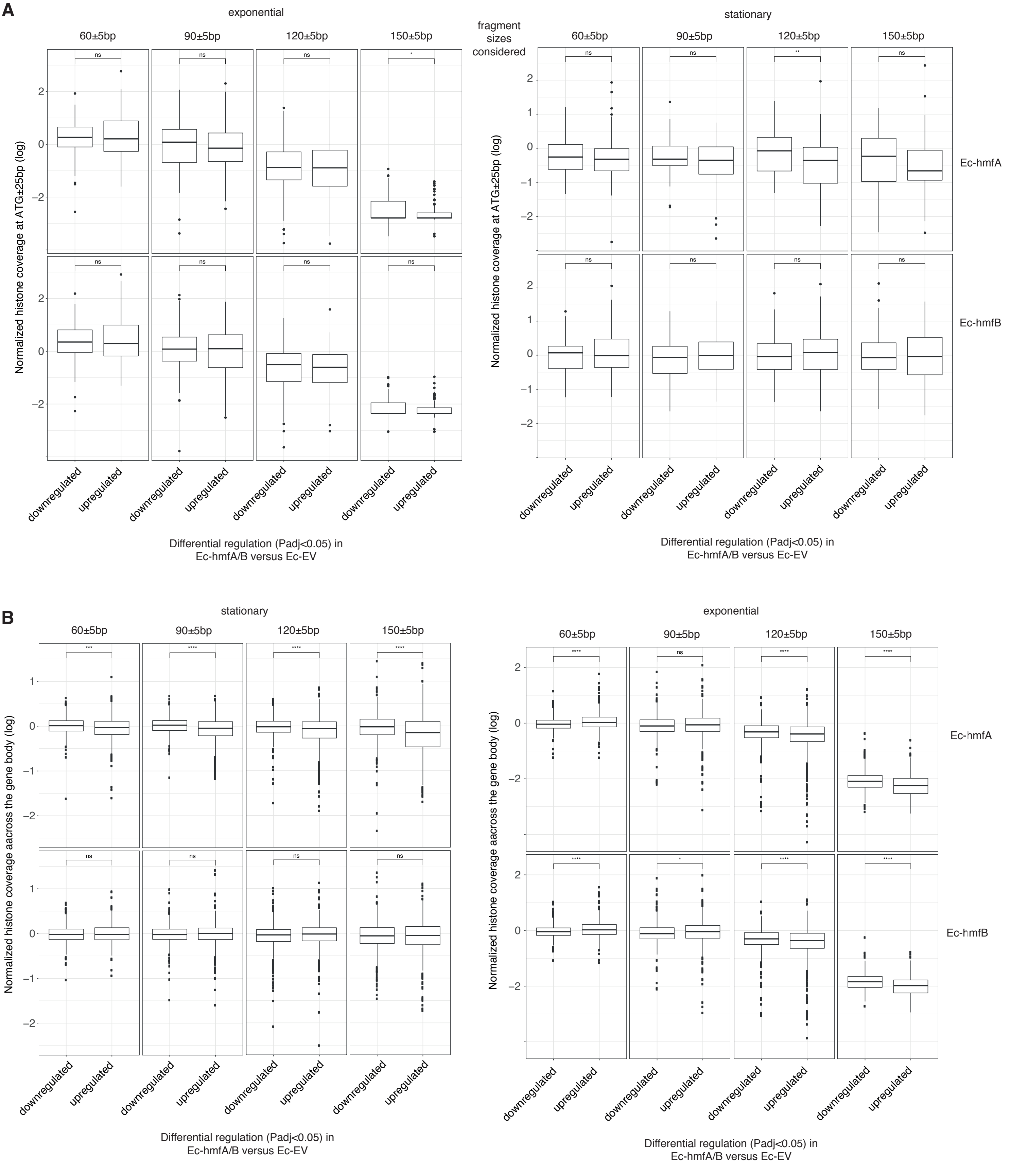

**Figure S4. The impact of archaeal histones in *E. coli* on transcription. A.** Reduced transcript abundance in histone-expressing strains is not associated with higher average histone occupancy around the ATG (±25bp). Only 5' genes in operons that are represented in Figure 4A are considered in this analysis, to enable a fair comparison between occupancy at transcription and translation start sites. **B.** Histone occupancy across the gene body for genes that are significantly up- or down-regulated in histone-expressing strains. ****P<0.001; ***P<0.005; **P<0.01; *P<0.05; expo: exponential phase; stat: stationary phase.

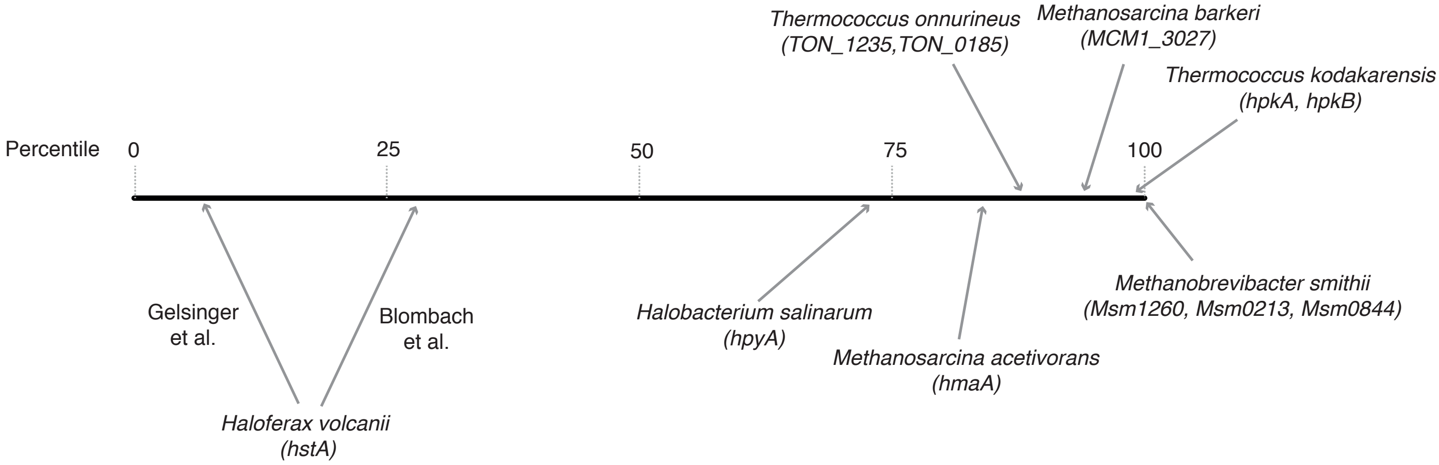

**Figure S5. Relative transcript levels of histone genes across different archaeal species.** Histones are assigned a percentile rank based on their relative expression in a given species and transcriptomic dataset (0=least abundant mRNA in the dataset; 100=most abundant mRNA in the dataset). For species with more than one histone gene, transcript levels were summed across histone genes. Because of signficant variability between studies, two separate estimates are given for *H. volcanii*. Data sources: *H. salinarum* (Gene Expression Omnibus accession GSE99730), *M. barkeri* (GSE70370), *T. onnurineus* (GSE85760), *M. acetivorans* (GSE64349), *M. smithii* (GSE25408), *H. volcanii* (Blombach et al. 2018 Nucl Acid Res 46:2308-2320; Gelsinger et al 2018 J Bacteriol 200:e00779-17), *T. kodakarensis* (Jäger et al. 2014 BMC Genomics 15:684).

**Table S1.** ***E. coli*** **K-12 MG1655-derived strains constructed for this study**

| Strain name | Plasmid name | Insert sequence (Histone CDS) |
|---|---|---|
| Ec-hmfA | pD681-hmfA | ATGGGCGAGCTGCCAATTGC<br>GCCGATCGGCCGCATTATCA<br>AAAATGCCGGTGCGGAGCGT<br>GTGAGCGACGACGCACGTAT<br>CGCGCTGGCAAAGGTTCTGG<br>AAGAAATGGGTGAAGAAAT<br>TGCCTCCGAAGCTGTCAAAT<br>TGGCAAAACACGCGGGTCGT<br>AAGACGATCAAAGCCGAAGA<br>TATCGAGCTGGCGCGCAAAA<br>TGTTTAAGTAA |
| Ec-hmfB | pD681-hmfB | ATGGAACTGCCAATTGCCCC<br>TATCGGTCGTATTATTAAAG<br>ACGCTGGTGCCGAGCGCGTG<br>AGCGATGACGCGCGCATCAC<br>CCTGGCAAAGATTCTGGAAG<br>AAATGGGCCGTGACATTGCG<br>TCCGAGGCCATCAAACTGGC<br>ACGTCACGCGGGTCGTAAGA<br>CGATCAAAGCTGAAGATATC<br>GAGCTGGCAGTTCGTCGCTT<br>CAAAAAGTGA |
| Ec-EV | pD681 | no insert |
| Ec-hmfA$_{nb}$ | pD681-hmfA$_{nb}$ | ATGGGCGAGCTGCCGATTGC<br>GCCGATTGGTCGTATTATCA<br>CCAACGCTGGCGCGGAGAGC<br>GTTTCCGACGACGCGCGCAT<br>TGCATTGGCAAAGGTCCTGG<br>AAGAAATGGGTGAAGAAATC<br>GCAAGCGAAGCCGTGAAACT<br>GGCGAAACACGCGGGTCGTA<br>AGAAAATCAAAGCTGAAGAT<br>ATCGAGCTGGCCCGTAAAAT<br>GTTCAAGTAA |
| Ec-hmfB$_{nb}$ | pD681-hmfB$_{nb}$ | ATGGAACTGCCGATTGCGCC<br>GATCGGCCGCATTATCACCG<br>ACGCGGGTGCCGAGAGCGTG<br>AGCGATGACGCACGCATCAC<br>GCTGGCGAAGATTCTGGAAG<br>AAATGGGCCGTGACATCGCG<br>TCCGAGGCCATTAAACTGGC<br>ACGTCACGCGGGTCGTAAAA<br>AGATCAAAGCTGAAGATATT<br>GAGTTGGCAGTTCGCCGTTT<br>CAAGAAATAA |

**Table S2. Fourier filtering parameters.**

| Sample | Growth Phase | Fragment size (bp) | PcVal | Thr (Zsc) | Number of peaks |
|---|---|---|---|---|---|
| Ec-hmfA | Exponential | 55 - 65 | 0.02 | 0.25 | 27049 |
| Ec-hmfA | Exponential | 85 - 95 | 0.015 | 0.25 | 18983 |
| Ec-hmfA | Exponential | 115 - 125 | 0.0125 | 0.25 | 15795 |
| Ec-hmfA | Exponential | 145 - 155 | 0.0075 | 0.25 | 7050 |
| Ec-hmfB | Exponential | 55 - 65 | 0.0225 | 0.25 | 28023 |
| Ec-hmfB | Exponential | 85 - 95 | 0.0175 | 0.25 | 25174 |
| Ec-hmfB | Exponential | 115 - 125 | 0.0125 | 0.25 | 15590 |
| Ec-hmfB | Exponential | 145 - 155 | 0.01 | 0.25 | 14363 |
| Ec-EV | Exponential | 55 - 65 | 0.015 | 0.25 | 21375 |
| Ec-EV | Exponential | 85 - 95 | 0.015 | 0.25 | 12923 |
| Ec-EV | Exponential | 115 - 125 | 0.01 | 0.25 | 9015 |
| Ec-EV | Exponential | 145 - 155 | 0.0075 | 0.25 | 2800 |
| Ec-hmfA | Stationary | 55 - 65 | 0.02 | 0.25 | 22865 |
| Ec-hmfA | Stationary | 85 - 95 | 0.0125 | 0.25 | 15287 |
| Ec-hmfA | Stationary | 115 - 125 | 0.01 | 0.25 | 10936 |
| Ec-hmfA | Stationary | 145 - 155 | 0.0075 | 0.25 | 7608 |
| Ec-hmfB | Stationary | 55 - 65 | 0.0225 | 0.25 | 17770 |
| Ec-hmfB | Stationary | 85 - 95 | 0.0175 | 0.25 | 19272 |
| Ec-hmfB | Stationary | 115 - 125 | 0.0175 | 0.25 | 18240 |
| Ec-hmfB | Stationary | 145 - 155 | 0.0175 | 0.25 | 18361 |
| Ec-EV | Stationary | 55 - 65 | 0.02 | 0.25 | 9811 |
| Ec-EV | Stationary | 85 - 95 | 0.0175 | 0.25 | 12575 |
| Ec-EV | Stationary | 115 - 125 | 0.015 | 0.25 | 14465 |
| Ec-EV | Stationary | 145 - 155 | 0.015 | 0.25 | 13274 |
| *M. fervidus* | Exponential | 55 - 65 | 0.02 | 0.25 | 6418 |
| *M. fervidus* | Exponential | 85 - 95 | 0.015 | 0.25 | 5176 |
| *M. fervidus* | Exponential | 115 - 125 | 0.01 | 0.25 | 3707 |
| *M. fervidus* | Exponential | 145 - 155 | 0.01 | 0.25 | 3576 |
| *M. fervidus* | Stationary | 55 - 65 | 0.0225 | 0.25 | 8768 |
| *M. fervidus* | Stationary | 85 - 95 | 0.0175 | 0.25 | 7848 |
| *M. fervidus* | Stationary | 115 - 125 | 0.0125 | 0.25 | 5208 |
| *M. fervidus* | Stationary | 145 - 155 | 0.01 | 0.25 | 3798 |

**Table S3. Differential Expression Analysis Outliers**

| DEA* pair | Growth phase | Replicates excluded** |
|---|---|---|
| Ec-hmfA vs Ec-EV | Exponential | Ec-EV 1 |
| Ec-hmfB vs Ec-EV | Exponential | Ec-EV 1 |
| Ec-hmfA$_{nb}$ vs Ec-EV | Exponential | / |
| Ec-hmfA$_{nb}$ vs Ec-EV | Exponential | / |
| Ec-hmfA vs Ec-hmfB | Exponential | Ec-EV 1 |
| Ec-hmfA vs Ec-hmfA$_{nb}$ | Exponential | Ec-hmfA 6 |
| Ec-hmfB vs Ec-hmfB$_{nb}$ | Exponential | / |
| Ec-hmfA vs Ec-EV | Stationary | Ec-hmfA 2 |
| Ec-hmfB vs Ec-EV | Stationary | Ec-hmfA 2 |
| Ec-hmfA$_{nb}$ vs Ec-EV | Stationary | / |
| Ec-hmfB$_{nb}$ vs Ec-EV | Stationary | / |
| Ec-hmfA vs Ec-hmfB | Stationary | Ec-hmfA 2 |
| Ec-hmfA vs Ec-hmfA$_{nb}$ | Stationary | Ec-hmfA 2 |
| Ec-hmfB vs Ec-hmfB$_{nb}$ | Stationary | Ec-hmfB 1 & Ec-hmfB$_{nb}$5 |

* DEA: Differential expression analysis
** see NCBI Gene Expression Omnibus accession number GSE127680 for sample IDs

### Table S4. qPCR primer sequences.

| Target | Forward Primer Sequence | Reverse Primer Sequence | Organism |
|---|---|---|---|
| asmA | GACAACCTCATCCCGCTTAAT | TGATAGGCCGGTTCATCAATAC | *E. coli* |
| clpX | GTTGAATGAACTGAGCGAAGAAG | ATCCACGCCTTCCAGATTAAA | *E. coli* |
| cmk | ACTCAGGAAGTGGCGAATG | CGGCAATCAGACCTGGTAAT | *E. coli* |
| cpoB | CCAACTCCAGCAACAACTTTC | CCACGACCTGATTCAGTTGATA | *E. coli* |
| dcyD | GTGCCGATACGCTGATTACT | TCTGCGGTTGTGCCAATA | *E. coli* |
| dusB | GGTTTGGGAAAGCGACAAATC | GCTGCATCTGCCATTTCTTTC | *E. coli* |
| elyC | CCACGCCTGAATGAAGGTAT | CCGCTGTACTCACCGTATTG | *E. coli* |
| fabF | GTTGTCTCCTGTCGGCAATA | CGTTGCATAGGCGCTAGTAT | *E. coli* |
| fabI | CCTCCGGTATCAAAGACTTCC | CGCAGAGTTACCCACATCTT | *E. coli* |
| fliA | CCAGGAAGAGCTGAATCTCAAA | ACCCAGTTTAGTGCGTAACC | *E. coli* |
| gadB | TGGGTGGTCAAATTGGTACTT | GGTAAGAGGCGTTCTGTACTTT | *E. coli* |
| gloB | CCATATTTCTCACCCACCATCA | TTGTGTCTCTTGTGGACCATAC | *E. coli* |
| gnd | CCTCGTCGCATCCTGTTAAT | CGAATAGTGTCCTGGAAGAAGG | *E. coli* |
| hdeA | GGCGTTATTCTTGGTGGTCT | GAAATCTTCACAGGTCCAGGAG | *E. coli* |
| hflC | CCGGGTCTGCATTTCAAGATA | CGACGATCAGGTCTTTCTTCTC | *E. coli* |
| ibpA | CCGCTTTACCGTTCTGCTAT | GTTCAACGTTATACGGAGGGTAG | *E. coli* |
| iscS | ACACAAAGCGGTACTGGATAC | CATCGCTGCTTCAAGTTCTTTC | *E. coli* |
| leuD | TGCGCTGGTGAAAGCTAAT | CGGAAGGCATCGATGGTAAA | *E. coli* |
| lptD | AAGTACACCACCACCAACTAC | TGCCACGACGATGCATATAA | *E. coli* |
| map | GCACTATGACTCCCGTGAAA | CTTTCATGGTGCGGATCTCT | *E. coli* |
| mepM | GCACAAGAAGATGAAGCCATTC | CGCCAGTGGAAACAACATATTC | *E. coli* |
| mnmG | CCAGGTATCCGGTCTTTCTAAC | ACACCAGCAGAATGGAGATG | *E. coli* |
| murA | GTCGGCGAAGACTGGATTAG | TGAACTGGGCCTGCATATC | *E. coli* |
| nagA | CTGGATGAAGTGCTACGTATGG | GCAGTCAGGTTGGCTACTTTA | *E. coli* |
| ompX | GCAAGCTCTGGTGACTACAA | CATAACCCACACCCACTACAC | *E. coli* |
| rrlH | ACAGGTGGTCAGGTAGAGAATA | CTACATATCAGCGTGCCTTCTC | *E. coli* |
| rrsA | TCGGAATTACTGGGCGTAAAG | GACTCAAGCTTGCCAGTATCA | *E. coli* |
| rsmA | GCAGGATGCGATGACCTTTA | CAACGGCGTGGAGATGTTAT | *E. coli* |
| sodB | CATCGAGTATCACTACGGCAAG | CACCTTCAGAGCTGCGAATAA | *E. coli* |
| talB | AGGCATCAACTGTAACCTGAC | GTCAAGAATACGGCCAACAAAC | *E. coli* |
| torD | TACAAACAGGACGAGCAAGAG | TCGAGATAGATCGCCAGATGA | *E. coli* |
| yafD | GACAACGCCAGAGTTAGTACAG | AAGGGTCATTACGCCAGAAG | *E. coli* |
| yaiE | CGACTGGCAGGTGTATGAAG | GCACAGATAAGAGGTGGGTTC | *E. coli* |
| yccA | CGGACCTATTCTGAACACCTATC | CAGAGCAGCAGAAGAACACTA | *E. coli* |
| yebG | TGAAGCCCTTTCGCTATGG | GCATCTTCCTCTTCGGATTCA | *E. coli* |
| yfjD | CACTATTGTTGGGATGCGTTTG | TCGGCAATACCTCAGCAAATA | *E. coli* |
| yifL | CCCGCCAAATGATTATCGAAAC | ATGACAACGGCATCATCTTCT | *E. coli* |
| zitB | GTATGGATGGTAGGCGAGAAG | CCATCAGGTAGTGTTGGATCTG | *E. coli* |
| ACT1 | GAGGTTGCTGCTTTGGTTATTG | ACCGACGATAGATGGGAAGA | *S. cervisiae* |
| RDN18-1 | TGCTGGCGATGGTTCATT | CTCCGGAATCGAACCCTTATTC | *S. cervisiae* |
| RPN2 | GGTGGTTCATTGTATGGTTTGG | GCCTGAAGTTCCGCTATTCT | *S. cervisiae* |
